# Deep cerebellar tFUS engages cortical circuits via convergent local and sensory-driven mechanisms

**DOI:** 10.1101/2025.11.15.688639

**Authors:** Anvar Sariev, Hongchae Baek, Dajung Jung, Hyungmin Kim, Alexandr Pak

## Abstract

Transcranial focused ultrasound (tFUS) enables non-invasive access to deep brain structures with high spatial precision, but its mechanisms of action remain elusive and under active investigation. The leading hypothesis posits that tFUS exerts its neuromodulatory effects through direct activation of neurons, while others attribute effects to indirect activation of auditory pathways. Here, we investigated the neuromodulatory effects of tFUS on the dentato– thalamo–cortical (DTC) pathway by targeting either the lateral cerebellar nucleus (LCN) or auditory cortex (AUD) in anesthetized rats. Electroencephalography (EEG) recordings from bilateral motor cortices and the contralateral auditory cortex revealed that tFUS reliably evoked cortical potentials, whereas sham stimulation produced no responses. Both LCN and AUD stimulation activated the auditory cortex, suggesting auditory pathway involvements. However, only LCN stimulation elicited early and significant event-related potentials and gamma-band activity in the contralateral motor cortex, consistent with DTC pathway engagement. We propose that LCN-targeted stimulation engages the DTC pathway through modulation of neuronal excitability, while concurrent auditory inputs account for global cortical activation. These results are consistent with a hybrid mechanism in which tFUS modulates neuronal dynamics, converging both direct and indirect components depending on the stimulation site. Proposed framework reconciles competing views of tFUS as either a direct or indirect modulator and clarifies how ultrasound can differentially influence cortical circuits depending on the stimulation target.

## Introduction

Modulation of brain activity is a critical approach in both clinical neurology and neuroscience research. Broadly, brain stimulation techniques are classified as either invasive or non-invasive. Invasive approaches such as deep brain stimulation (DBS) have demonstrated therapeutic efficacy in disorders, such as Parkinson’s disease and essential tremor, but they carry inherent surgical risks and high procedural costs^1,2^. Non-invasive brain stimulation (NIBS) methods, including transcranial magnetic stimulation (TMS) and transcranial direct current stimulation (tDCS), offer safer alternatives but are often constrained by limited spatial resolution and depth of penetration^3,4^.

Transcranial focused ultrasound (tFUS) has recently emerged as a novel NIBS technique with the unique ability to deliver highly focused energy to both superficial and deep brain structures with millimeter precision, making it particularly promising for modulating subcortical pathways^5^. Several studies have reported the ability of tFUS to trigger neural activity in vitro ^6^, in vivo^7,8^, and in human ^9-11^. Although these findings demonstrate that tFUS can elicit measurable neural and behavioral effects, the exact mechanism has yet to be fully established, with the leading view suggesting that its effects are mediated through activation of mechanosensitive ion channels and cell-type specific voltage-gated channels^12-15^.

Several findings have questioned the neuromodulatory effect of tFUS, by claiming that neural activation during ultrasound stimulation is mainly due to cochlear activation of the auditory pathway ^16,17^. Furthermore, a very recent study in humans demonstrated that motor inhibitory effects previously attributed to tFUS stimulation were replicated but occurred equally during stimulation of a control (non-targeted) brain region and with sound presentation alone. This indicates that the observed inhibition was driven by auditory confounds rather than direct ultrasonic neuromodulation^18^. This uncertainty emphasizes the need for careful experimental validation to distinguish direct neural effects from auditory confounds and to clarify the true mechanisms of tFUS before its broader clinical application.

In this context, the dentato-thalamo-cortical (DTC) pathway represents an ideal target for dissecting both local and network-level effects of tFUS. The lateral cerebellar nucleus (LCN), a major output node of the cerebellum, projects to motor and non-motor cortical regions and plays a vital role in motor coordination and cognitive processing ^19,20^. Targeting the LCN with tFUS may offer a powerful non-invasive method for modulating broad cortical regions via this well-characterized tract – an approach that holds significant promise for applications such as post-stroke motor rehabilitation^21-25^.

In this study, we compared the effects of tFUS targeted to the LCN and the auditory cortex in healthy rats by measuring electroencephalographic (EEG) responses over motor and auditory cortices. Stimulation of the LCN, part of the DTC pathway, evoked robust, lateralized cortical responses in motor and auditory areas, with particularly stronger and earlier potentials observed in the contralateral motor cortex—suggesting pathway-specific engagement. In contrast, stimulation of the auditory cortex did not produce significant lateralized responses in motor cortex. These findings indicate that tFUS can induce both localized and network-level modulatory effects, with cerebellar stimulation showing distinct long-range activation patterns, thereby reinforcing its potential for therapeutic applications such as stroke rehabilitation.

## Methods

### Animal Preparation

The study was conducted on eight healthy Sprague Dawley rats (250-280 g, 8-10 weeks old). The animals were housed in a temperature-controlled room with a 12h light/dark cycle and had ad libitum access to water and food. All surgical procedures were performed under anesthesia induced by ketamine (80 mg/kg) and xylazine (10 mg/kg). The experimental protocol was reviewed and approved by the Institutional Animal Care and Use Committee of the Korea Institute of Science and Technology.

### Sonication setup

A 0.35 MHz single-element concave-shaped focused ultrasound transducer (GPS350-D25_FL25, The Ultran Group, USA) with an aperture size of 25 mm, a focal length of 23.5 mm, and lateral and axial full-width at half-maximum (FWHM) dimensions of 5 mm and 29 mm, respectively, was employed in our experiments. The transducer’s FWHM dimensions, focal length, and intensity were measured in degassed and deionized water using a calibrated hydrophone (HNR-0500, ONDA Corp., CA, USA). The experimental setup consisted of two function generators (33210A, Agilent, Santa Clara, CA), an oscilloscope (DPO4104, Tektronix Inc. USA), and a linear radio-frequency power amplifier (240L, ENI Inc., Rochester, NY).

The tFUS exposure conditions used in our experiments were as follows: 50 % of duty cycle, 1 kHz of pulse-repetition frequency (PRF), 0.5 ms of pulse duration (PD), 300 ms of pulse train duration (PTD), 3 s of pulse train repetition interval (PTRI), and a spatial-peak pulse-average intensity (I_sppa_) of 2.54 W/cm^2 26^. This particular sonication parameter has been proven to be effective for the rehabilitation of ischemic stroke in mouse models in the previous studies^24,27^. During stimulation sessions transducer was coupled with ultrasound gel and focused on the LCN, a key node in the DTC pathway, or the primary auditory cortex (AUD) based on pre-marked skull targets (LCN: AP -11.5 mm, ML -3.5 mm, DV: -6.0 mm; Au1: AP -5.5 mm, ML -7 mm, DV -3 mm from bregma) with a customized transducer holder^25^. The sham session was conducted with the transducer uncoupled from the skull surface to exclude any potential effects of audible sound generated by the transducer.

### Electrode implantation and electrophysiological recordings

Electrode implantation and recording were performed on anesthetized animals fixed on a stereotaxic frame. Anesthesia was maintained below motor response threshold to toe pinch (areflexic state). The skin over the skull was exposed, and small burr holes were drilled over both the primary motor cortex (M1, AP: 2.5 mm, and ML: ±3 mm from bregma) and the contralateral to the stimulation site primary auditory cortex (Au1, AP: -5.5 mm, and ML: -7 mm from bregma). Small screws (Antrin Miniature Specialties, AMS120/1B-25) soldered to subdermal electrode wires (SWE-L25, Ives EEG Solution, MA, USA) were fixed into the holes.

Three-channel EEG signals were acquired using a data acquisition system (PL3508, Model ML138, ADInstruments, Sydney, Australia) and digitized at a 1 kHz sampling rate, band-pass filtered (0.3-200 Hz), and notch-filtered (60 Hz) using LabChart8 software (ADInstruments, Sydney, Australia). Cortical EEG data were acquired during stimulation and sham sessions.

### Electrophysiological data analysis

EEG data were processed using LabChart8 software (ADInstruments, Sydney, Australia) and custom scripts written in Matlab (The MathWorks, Inc., Natick, MA) and Python. Data quality was assessed using the baseline (−200 to 0 ms) root-mean-square (RMS) voltage. Sessions were excluded if their RMS exceeded the cohort median ± 2.5 standard deviations within the animal, ensuring that only recordings with stable baseline noise levels were included in further analysis. One animal was excluded from the auditory tFUS condition due to elevated baseline RMS. Consequently, six rats were included in the LCN and SHAM conditions, and five rats in the auditory stimulation condition. To directly compare ERPs during LCN or auditory cortex stimulation versus SHAM conditions, we employed cluster-based permutation testing using the MNE-Python software package^28^. This non-parametric approach effectively controls for multiple comparisons across time points while maintaining statistical power to detect genuine effects^29^. For ERP amplitude comparisons, we used one-dimensional cluster-based permutation tests with the following parameters: number of permutations = 1000, cluster-forming threshold = 10. The analysis was performed separately for each electrode (contralateral M1, ipsilateral M1, and auditory cortex) over a time window of -200 to 1000 ms relative to stimulation onset. Statistical significance was assessed at α = 0.05, with cluster-level correction for multiple comparisons. Only clusters surviving the permutation-based correction were considered statistically significant.

#### Peak-to-Peak Amplitude Analysis

In addition to cluster-based permutation testing, we performed targeted analysis of early response components by identifying peak positivity and negativity within the first 50 ms following stimulation onset. Peak-to-peak amplitudes were calculated as the difference between the maximum positive and negative deflections within this early response period. To assess the significance of stimulus-evoked responses, peak-to-peak amplitudes were compared to baseline activity using the Wilcoxon signed-rank test. Baseline activity was characterized using the mean amplitude during the pre-stimulation period (-200 to 0 ms).

#### Time-Frequency Analysis

Time-frequency decomposition was performed using complex Morlet wavelet convolution, adapted from established methods^30,31^. The analysis computed multiple measures of neural oscillatory activity including total power, induced (non-phase-locked) power, evoked (phase-locked) power, and inter-trial phase coherence (ITPC). Wavelets were applied across a frequency range of 2-50 Hz using 40 logarithmically-spaced frequencies, with wavelet cycles ranging from 3 to 10 (logarithmically increasing with frequency) to optimize the trade-off between temporal and frequency resolution. Evoked power measured phase-locked activity from the ERP spectral content, and inter-trial phase coherence (ITPC) quantified phase consistency across trials, calculated as the absolute value of the mean unit vector - *mean*(*e*^*iφ*^), where φ represents the instantaneous phase. All power measures were baseline-corrected using the pre-stimulation period (-200 to 0 ms) and expressed in decibels relative to baseline using the formula 10*log_10_(power/baseline_power). ITPC values range from 0 (random phase) to 1 (perfect phase consistency). We computed mean spectral power and ITPC values within four canonical frequency bands during the 200 ms period following stimulation onset: theta (4-10 Hz), alpha (8-12 Hz), beta (13-30 Hz), and gamma (30-50 Hz). For each frequency band, power and ITPC values were averaged across the specified frequency range and the 200 ms post-stimulus time window for each trial, then averaged across trials to obtain single values per condition and electrode.

Statistical analysis of spectral power between conditions was conducted using MNE cluster-based permutation tests with 1,000 permutations and a cluster-forming threshold of t = 10, in order to identify significant differences between stimulation conditions (LCN vs. sham, AUD vs. sham) and between hemispheres (contralateral vs. ipsilateral motor cortex). For ITPC comparisons, we implemented a custom cluster-based permutation test specifically designed for phase coherence data. This analysis used 1,000 permutations with α = 0.05 for final cluster significance and a cluster formation threshold of α = 0.05. The algorithm calculated ITPC for each condition, computed t-statistics for group differences, identified clusters using statistical thresholds based on degrees of freedom (df = n_1_ + n_2_ - 2), and determined cluster significance by comparing observed cluster masses (the sum of t-values within clusters) to a null distribution generated through trial-label permutation. Only clusters with a minimum size of ≥ 5 time-frequency points were considered for analysis. This approach provides robust statistical inference for phase coherence differences while controlling for multiple comparisons across the time-frequency domain and properly handling the circular nature of phase data. Between-hemisphere comparisons (contralateral vs ipsilateral motor cortex) for each frequency band were performed using the Wilcoxon signed-rank tests.

## Results

### LCN-targeted tFUS evokes lateralized cortical responses

To investigate the effects of tFUS on brain dynamics, we applied sonication to the deep lateral cerebellar nucleus (LCN) or primary auditory cortex (AUD) of anesthetized rats and recorded evoked cortical activity using EEG electrodes placed over the bilateral primary motor cortices (M1-ipsi and M1-contra) and the contralateral auditory cortex (Au1). Each recording session included baseline, sham, and stimulation blocks, with sham stimulation serving as a control to account for potential non-specific sensory effects (Fig. 1A).

**Figure 1.**
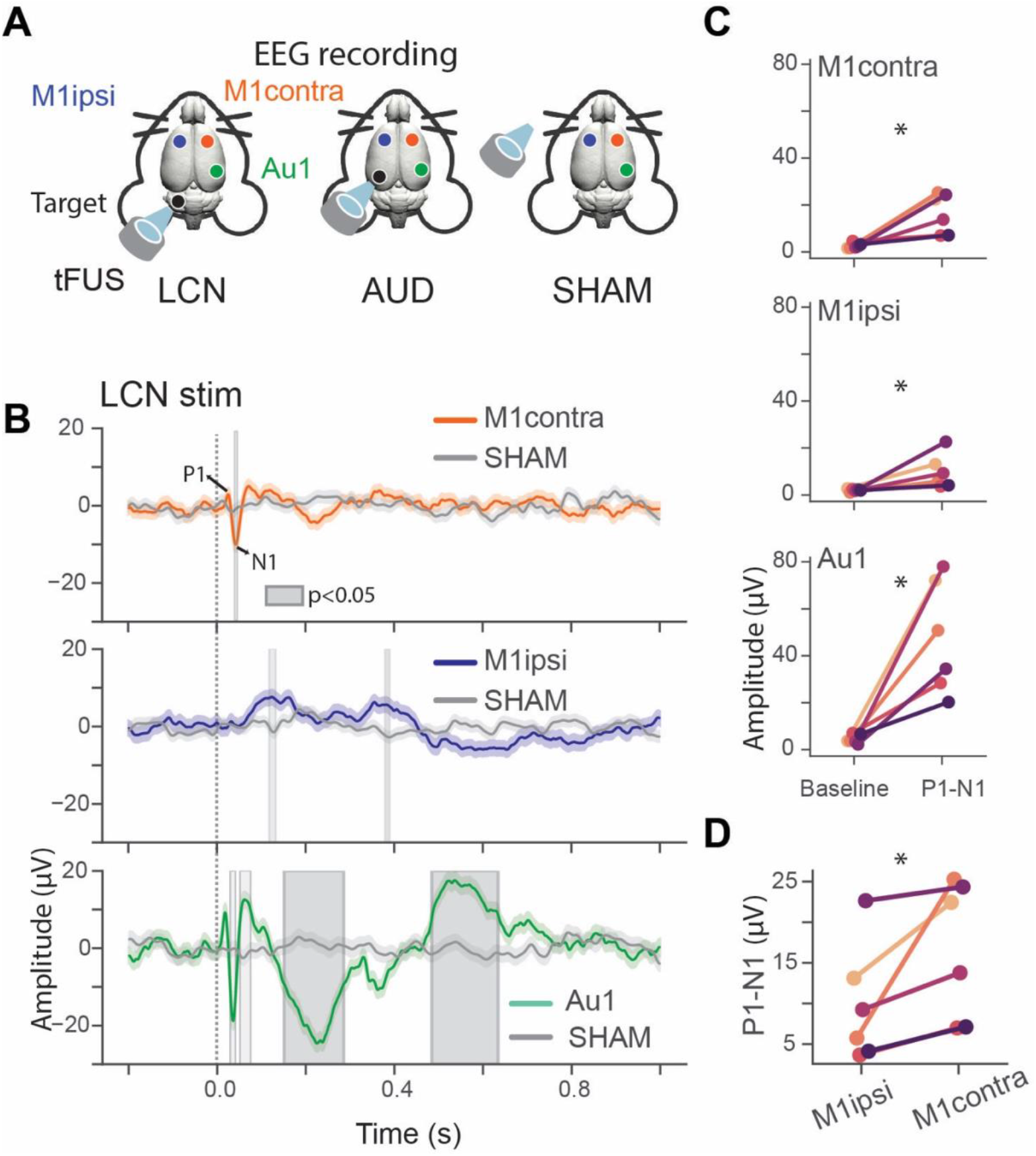
tFUS targeting the LCN evokes robust and lateralized cortical responses. **A)** Experimental setup showing screw electrode placement over bilateral motor cortices (M1) and auditory cortex (Au1), and transducer positioning for LCN or AUD stimulation. Bottom: Experimental protocol depicting baseline, sham (transducer uncoupled), and active stimulation blocks. **B)** Average EEG traces during LCN stimulation overlaid with sham stimulation (gray) recorded from contralateral M1 (orange), ipsilateral M1 (blue), and auditory cortex (green) (n=6 rats, LCN: 1228 trials, SHAM: 1119 trials). Shaded gray regions indicate statistically significant differences between LCN and SHAM conditions based on cluster-based permutation tests (p<0.05). **C)** Peak-to-peak amplitude (P1-N1) comparison between post-stimulation (0-100 ms) and mean baseline (-200 to 0 ms) periods across recording channels. Each dot represents one rat with connecting lines showing paired measurements. (Wilcoxon signed-rank test; M1-contra vs. baseline: W=0, p = 0.03; M1-ipsi vs. baseline: W= 0, p = 0.03; Au1 vs. baseline: W=0, p = 0.03) **D)** Direct comparison of P1-N1 amplitudes between contralateral and ipsilateral M1 during LCN stimulation. Each dot represents one rat with connecting lines showing paired measurements (Wilcoxon signed-rank test; M1-contra vs M1-ipsi: W=0, p = 0.03).

Stimulation of the LCN reliably evoked cortical responses in both motor and auditory cortical areas compared to SHAM (Fig. 1B, cluster-based permutation tests, p<0.05). At the contralateral motor cortex, clear event-related potentials (ERPs) were observed, characterized by distinct P1 and N1 components occurring within the first 100 ms following stimulus onset. In contrast, ERPs recorded from the ipsilateral motor cortex exhibited slower temporal dynamics, with broad positive deflections peaking around 150 ms and again near 400 ms post-stimulation. Additionally, the auditory cortex displayed more complex and prolonged ERP waveforms, with multiple peaks extending beyond 400 ms post-stimulation (cluster-based permutation tests, p<0.05).

To assess the strength of cortical responses, we compared the P1-N1 peak-to-peak amplitudes during LCN stimulation with baseline activity across all recording sites (Fig. 1C). M1-contra channel exhibited a mean P1-N1 amplitude of 28.2 ± 6.1 μV. In contrast, the M1-ipsi channel showed a lower amplitude of 15.5 ± 5.3 μV—the weakest response among the recorded regions. Notably, the Au1 channel displayed the strongest activation, with a P1-N1 amplitude of 49.3 ± 4.9 μV. These effects were consistent across animals, and statistical comparisons confirmed significant increases in ERP amplitude relative to baseline at all sites (Wilcoxon signed-rank test, *W* = 0, *p* = 0.03). A direct comparison of P1-N1 amplitudes between the motor cortices revealed significantly greater responses in the contralateral M1 than in the ipsilateral side (Fig. 1D; Wilcoxon signed-rank test, *W* = 0, p = 0.03). This lateralized response supports the involvement of the contralateral DTC pathway in mediating the cortical effects of tFUS.

These findings demonstrate that focused ultrasound stimulation of the LCN produces significant, time-locked cortical activity, with large and early ERPs in the contralateral motor cortex, consistent with the engagement of the DTC pathway. The concurrent activation of the auditory cortex suggests additional network-level effects or multisensory integration. The absence of significant ERP responses during sham stimulation supports the exclusion of non-specific sensory confounds.

### LCN stimulation induces lateralized gamma-band synchronization in the motor cortex

To investigate the spectral and temporal features of cortical responses to LCN stimulation, we performed time-frequency analysis on EEG signals. Time-frequency spectrograms revealed a robust increase in evoked power primarily in gamma band (30–50 Hz) within the first 200 ms post-stimulation in the contralateral M1. This gamma-band response was substantially reduced in the ipsilateral M1, indicating hemispheric asymmetry in stimulus-evoked oscillatory activity. The auditory cortex also exhibited early gamma-power increases, along with prolonged low-frequency activity in the alpha and theta bands, suggesting broad cortical engagement (Fig. 2B).

**Figure 2.**
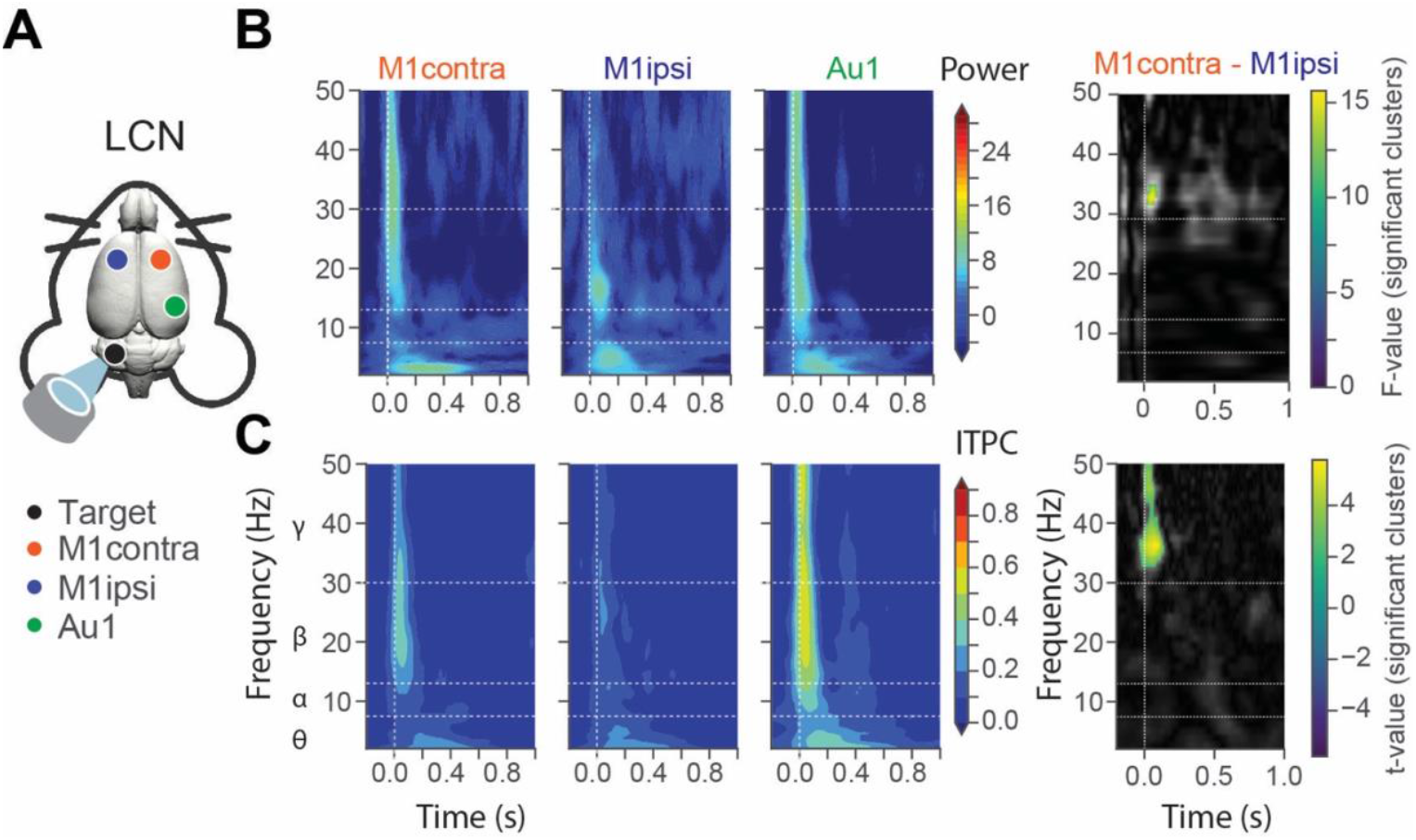
Time-frequency analysis of EEG traces reveals hemisphere-specific oscillatory responses to tFUS stimulation of LCN. A) Schematic representation of experiment with coupled transducer focused on LCN. B) Time-frequency spectrograms showing evoked power changes in contralateral M1, ipsilateral M1, and auditory cortex during LCN stimulation (left). Statistical comparison heatmap of difference between contralateral and ipsilateral M1 showing F-values from cluster-based permutation tests (right, colored regions – statistically significant values, black/white – not significant values). C) ITPC maps demonstrating hemispheric asymmetry in phase locking. Statistical comparison heatmap of difference between contralateral and ipsilateral M1 showing t-values from cluster-based permutation tests (right, colored regions indicate statistically significant clusters; black/white regions indicate non-significant differences).

Inter-trial phase coherence (ITPC) analysis revealed similarly lateralized effects. Strong phase locking in the gamma band was observed in the M1-contra and Au1 during the early post-stimulus period (0–200 ms), while the ipsilateral M1 showed markedly weaker phase consistency (Fig. 2C). Cluster-based permutation testing confirmed that these hemispheric differences were statistically significant. Specifically, comparisons of power between M1-contra and M1-ipsi revealed significant clusters in the gamma band (*p* < 0.05), while ITPC analysis also showed significant clusters in the same frequency range (Fig. 2B and 2C, right panel). Average spectral power within a 0–0.2 s time window post-stimulation in the gamma band (Fig. S1) was significantly higher in the contralateral motor cortex compared to the ipsilateral motor cortex (Wilcoxon signed-rank test, *W* = 0, *p* = 0.03). Also, comparison of average ITCP values showed borderline significance in gamma band (Wilcoxon signed-rank test, *W* = 0, *p* = 0.06).

These results demonstrate that LCN stimulation elicits rapid and coherent gamma-band activity in the contralateral motor cortex, consistent with activation of the DTC pathway. The lateralized increase in both spectral power and phase-locking further supports the specificity of tFUS effects and highlights its utility in probing functional connectivity in targeted neural circuits.

### Auditory cortex targeted tFUS produces bilaterally symmetric cortical responses

To evaluate whether tFUS applied to a cortical target produces modulatory effects similar to those observed with deep cerebellar stimulation, we delivered tFUS to the left primary auditory cortex (AUD), while recording with same cortical channels. In contrast to the pronounced and lateralized responses observed during LCN stimulation (Fig. 1), auditory cortex stimulation evoked only modest EEG deflections in motor cortices, with comparable temporal profiles across hemispheres. The ipsilateral M1 exhibited broad peaks at approximately 200 and 1000 ms after stimulation, whereas the contralateral M1 did not show any significant peaks. The Au1 channel displayed a broad and prolonged ERP response, mainly between 100 to 400 ms, compared to sham stimulation (cluster-based permutation test, p<0.05).

Peak-to-peak (P1–N1) amplitude analysis confirmed the absence of robust evoked responses (Fig. 3B). Although all three channels showed a mean amplitude increase, this did not reach significance compared to baseline (Wilcoxon signed-rank test, M1-contra vs. baseline (W=1, p=0.125), M1-ipsi vs baseline (W=0, p=0.06), Au1 vs baseline (W=0, p=0.06)). Direct comparison of contralateral versus ipsilateral M1 amplitudes during AUD stimulation (Fig. 3C) revealed no hemispheric asymmetry (W = 3, p = 0.31).

**Figure 3.**
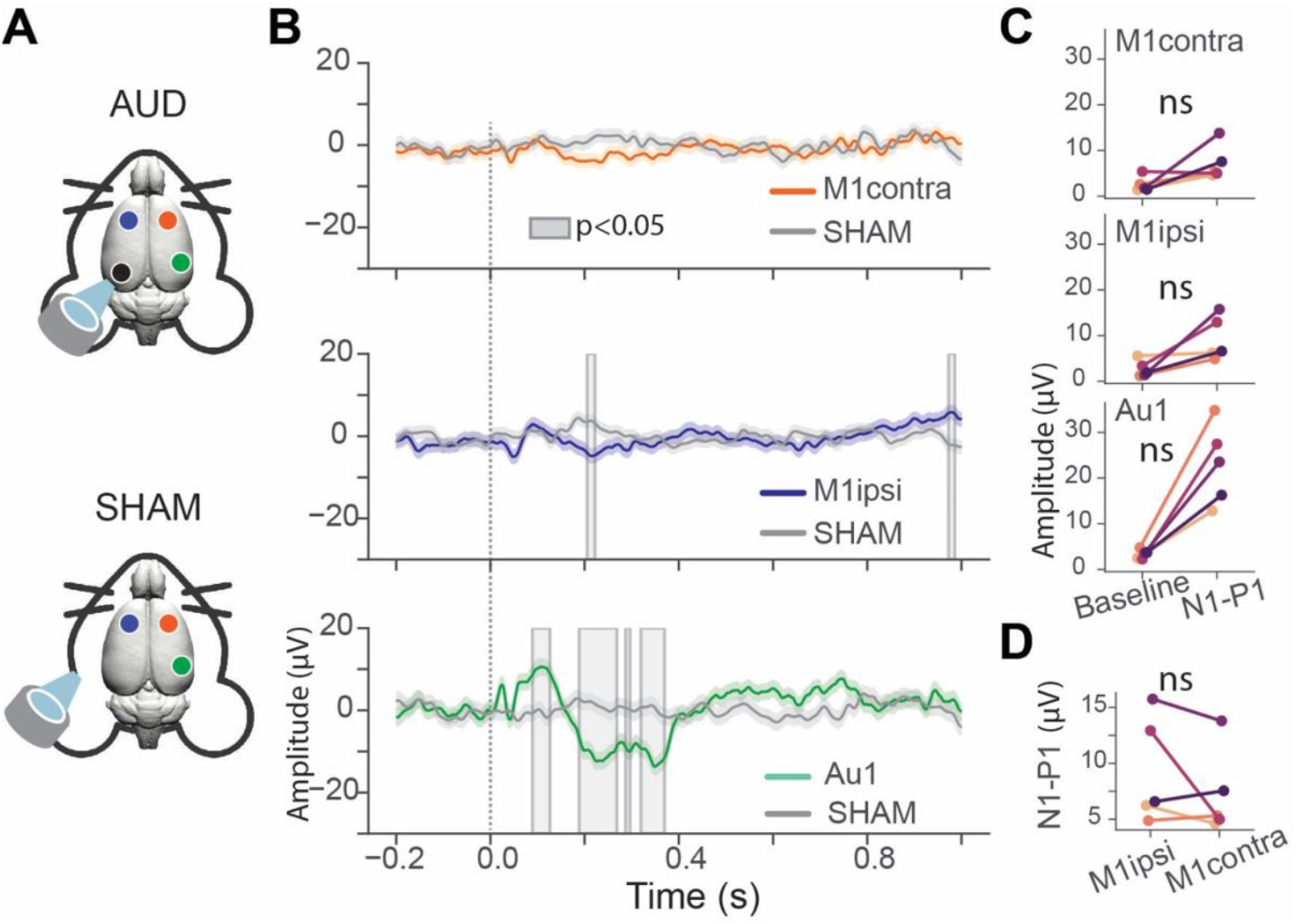
tFUS stimulation of auditory cortex produces similar responses in both hemispheres. **A)** Average EEG traces during AUD stimulation overlaid with sham stimulation (gray lines) recorded from contralateral M1 (orange), ipsilateral M1 (blue), and contralateral auditory cortex (green) (n=5 rats, AUD: 990 trials, SHAM: 1119 trials). Shaded gray regions indicate statistically significant differences between AUD and SHAM conditions using cluster-based permutation tests (p<0.05). **B)** Peak-to-peak amplitude (P1-N1) comparison between post-stimulation (0-100 ms) and mean baseline (-200 to 0 ms) periods across recording channels. Each dot represents one rat with connecting lines showing paired measurements (Wilcoxon signed-rank tests: M1-contra vs baseline, W=1, p=0.125; M1-ipsi vs baseline, W=0, p=0.06; Au1 vs baseline; W=0, p=0.06). No significant differences observed. **C)** Direct comparison of P1-N1 amplitudes between contralateral and ipsilateral M1 during AUD stimulation. Each dot represents one rat with connecting lines showing paired measurements (Wilcoxon signed-rank test: M1-contra vs M1-ipsi; W=3, p=0.31).

When compared with LCN stimulation results, which elicited large, statistically significant, and hemispherically asymmetric ERPs—particularly in the contralateral motor cortex—auditory cortex stimulation produced smaller, statistically non-significant responses without lateralization. This suggests that targeting a cortical auditory area with tFUS engages local circuits but does not recruit the DTC pathway in the same manner as deep cerebellar stimulation.

### Auditory cortex stimulation evokes bilateral low-frequency activity without lateralization

To examine the spectral and temporal features of cortical responses to AUD stimulation, we performed time–frequency analysis on EEG signals (Fig. 4A). In contrast to the robust and asymmetric changes observed during LCN stimulation, spectrograms revealed brief and low-amplitude power increases across both M1 channels, with the most prominent response observed in Au1. These changes were confined to the first 200 ms post-stimulation and lacked sustained oscillatory activity (Fig. 4B).

**Figure 4.**
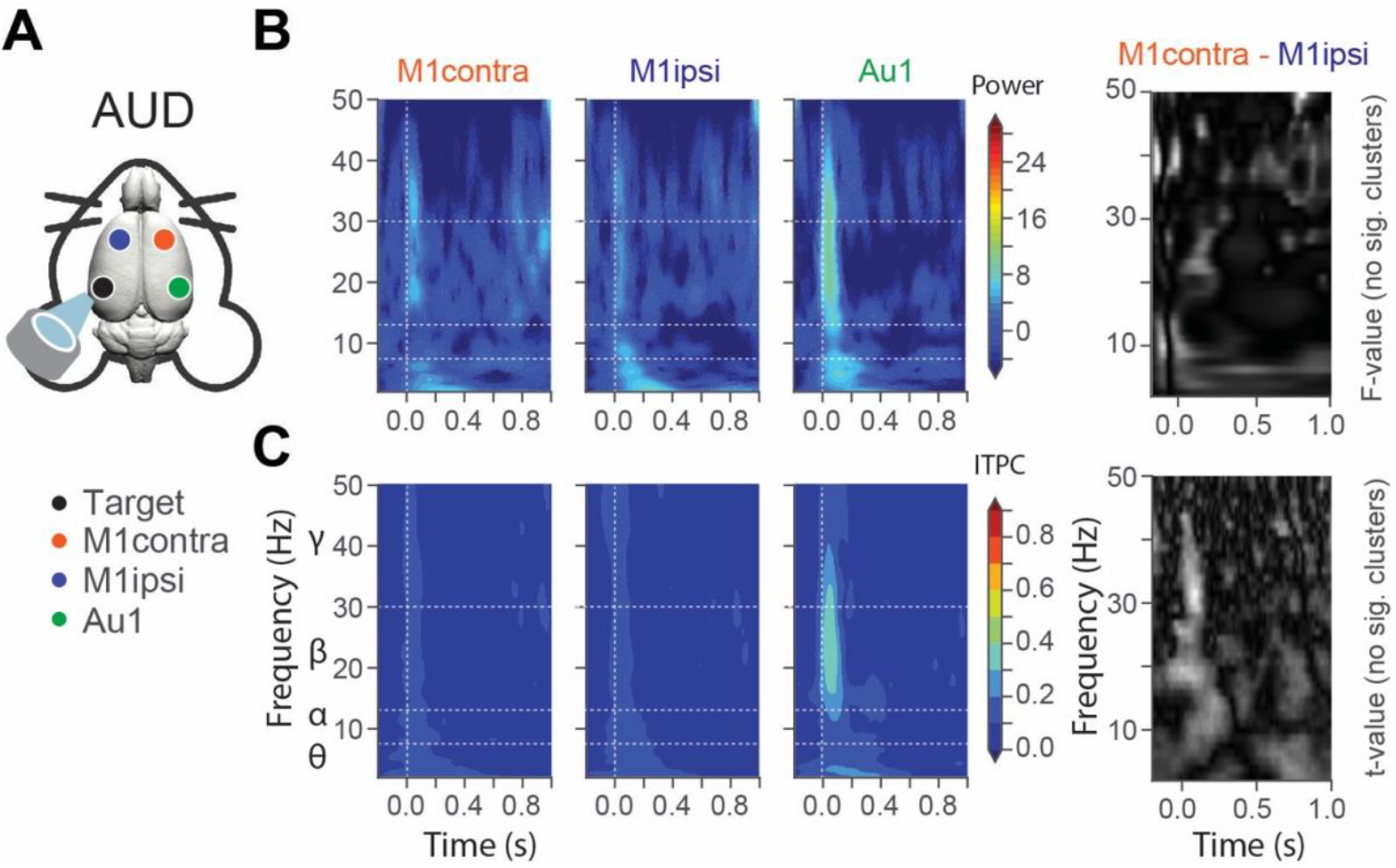
Time-frequency analysis reveals equivalent responses to tFUS stimulation of auditory cortex across hemispheres. A) Schematic representation of experiment with coupled transducer focused on auditory cortex (AUD). B) Time-frequency spectrograms showing evoked power changes in contralateral M1, ipsilateral M1, and auditory cortex during AUD stimulation (left). Statistical comparison heatmap of difference between contralateral and ipsilateral M1 showing F-values from cluster-based permutation tests (right, black/white regions – statistically not significant values). C) ITPC maps during AUD stimulation for the same channels demonstrating no hemispheric asymmetry in phase locking (left). Statistical comparison heatmap of difference between contralateral and ipsilateral M1 showing t-values from cluster-based permutation tests (right, black/white regions indicate non-significant differences).

ITPC analysis showed transient phase alignment in low- and mid-frequency ranges within Au1, while both motor cortex channels exhibited very weak or absent phase-locking responses (Fig. 4C). Cluster-based permutation testing revealed no significant differences between M1-contra and M1-ipsi in either power or ITPC across the 0–0.2 s post-stimulation window (p > 0.05 for all clusters; Fig. 4C).

Average spectral power was compared between M1 channels across canonical frequency bands (theta: 4–10 Hz, alpha: 8–12 Hz, beta: 12–30 Hz, gamma: 30–50 Hz) within the early post-stimulation period. No significant interhemispheric differences were detected in any band. Similarly, mean ITPC values across the same frequency ranges were equivalent between hemispheres (Fig. S2; Wilcoxon signed-rank tests, all p > 0.05).

These results demonstrate that AUD stimulation elicits weak and bilaterally symmetric oscillatory responses, with no evidence of hemispheric bias in either spectral power or phase-locking. The absence of lateralization stands in sharp contrast to the gamma-band–dominated, hemisphere-specific responses observed with LCN stimulation, suggesting that cortical targeting primarily engages local networks without strongly recruiting long-range subcortical pathways.

### Proposed framework for tFUS-induced modulation

The ongoing debate on how tFUS affects brain activity centers on whether its effects arise from local neuromodulation or from indirect sensory activation, particularly through the auditory pathway. Integrating previous findings with our current data, we outline three conceptual models that might explain how tFUS applied to LCN generates the cortical activation patterns observed in our experiments (Fig. 5).

**Figure 5.**
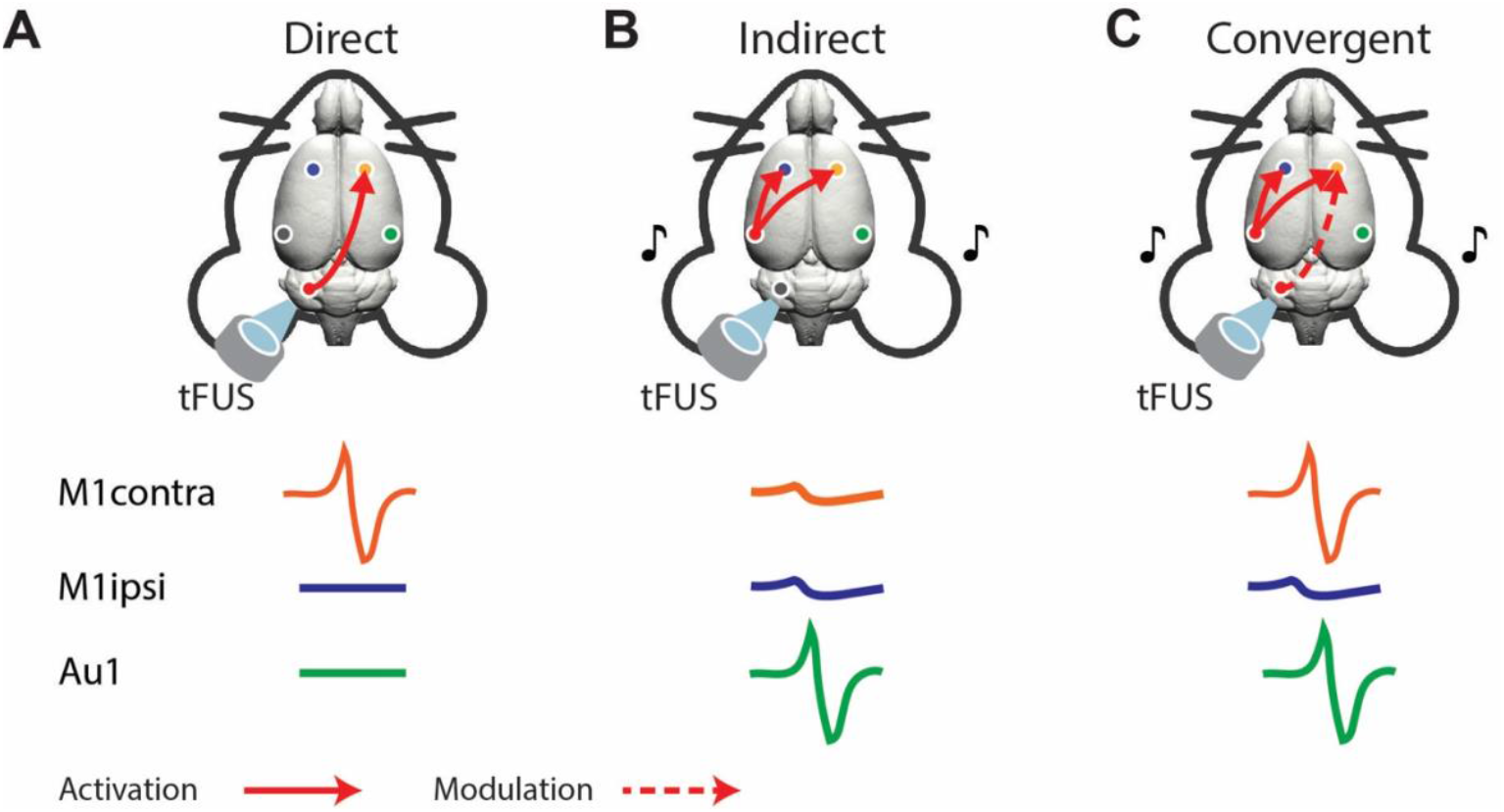
Proposed mechanisms of cortical activation during LCN stimulation. A) Direct activation model: tFUS excites LCN neurons, driving cortical responses via the DTC pathway. B) Indirect activation model: Cortical responses arise solely from cochlear activation of the auditory pathway. C) Convergent model (proposed): tFUS modulates LCN neuronal excitability without directly eliciting spikes, lowering the threshold for activation. Concurrent cochlear activation provides excitatory input, producing stronger contralateral M1 responses via cerebellar–cortical connections. This model may reconcile prior conflicting reports and explains the observed M1 asymmetry with LCN but not AUD stimulation.

#### Direct target activation

tFUS directly excites neurons within the focused target (Fig. 5A). This model could explain the contralateral M1 enhancement we observed in our experiment but does not explain off target activation of auditory cortex. Also, it fails to explain studies showing no tFUS-induced cortical activation in deaf mouse models^16,17,32^. *Indirect activation*. Cortical responses result primarily from acoustic coupling of ultrasound into the cochlea (Fig. 5B), driving the auditory cortex and associated cortical regions via ascending auditory pathways^33,34^. This mechanism aligns with our observation of auditory cortex activation in both LCN and AUD stimulation but fails to account for the lateralized M1 response detected during LCN stimulation. *Convergent sensory–modulatory mechanism*. In this framework, tFUS acts as a subthreshold modulator of neuronal excitability at the target while cochlear activation provides the dominant excitatory drive (Fig. 5C). In this view, ultrasound exposure at the LCN lowers the firing threshold of neurons within the DTC pathway, enabling sensory-driven input— originating from cochlear activation—to more effectively recruit contralateral motor cortical circuits.

This combined mechanism may reconcile previous seemingly opposing results: studies emphasizing direct local excitation and those attributing tFUS effects solely to auditory artifacts. In both our LCN and auditory cortex stimulation conditions, robust auditory cortex activation was observed, consistent with a cochlear contribution. However, only LCN stimulation produced asymmetric activation of contralateral M1, indicating pathway-specific modulation of cerebellar–cortical connectivity. These results are consistent with that tFUS primarily biases the excitability of target circuits, enabling sensory inputs to drive selective network activation— a framework that integrates both target-specific modulation and sensory-mediated excitation within a unified mechanistic model.

## Discussion

We show that tFUS produces two concurrent effects in the intact rodent brain. First, both LCN and AUD stimulation elicited robust activation of the auditory cortex, consistent with indirect cochlear-driven responses. Second, LCN stimulation—but not AUD stimulation—produced earlier and larger ERPs in contralateral M1 with lateralized increases in spectral power and inter-trial phase coherence, indicating pathway-specific engagement of the DTC circuit. Although the motor response was weaker than the auditory response, the combination of hemispheric asymmetry and time-frequency signatures provides strong evidence for pathway specific neural modulation.

The tFUS field has been split between reports emphasizing direct neural modulation at the target and those attributing responses to indirect auditory pathway activation ^35-38^. Several studies showing focal modulation of cortical and thalamic responses in humans and animals supports a direct component ^7,9,10,39,40^, whereas other studies have demonstrated that ultrasound can strongly drive the cochlea, producing widespread cortical activity that can dominate the readout ^16,17,32,41^. Our data accommodate both views: we consistently observe auditory cortex activation (supporting the indirect pathway), yet also detect a smaller, lateralized motor response only when stimulating LCN, in line with the known contralateral projections from the LCN through motor thalamus to M1, providing a circuit-level substrate for the observed asymmetry^42^.

In our study time–frequency analyses provide the clearest support for pathway specificity. During LCN stimulation, contralateral M1 exhibited enhanced gamma-band power with reliable phase-locking, while ipsilateral M1 showed weak or absent phase consistency (Fig. 2). Such higher-frequency, phase-coherent activity is associated with motor planning/execution and selective engagement of motor control networks^43,44^. In contrast, AUD stimulation showed weak bilaterally symmetric spectral and phase patterns in motor cortices (Fig. 4), consistent with generalized cortical activation from cochlear drive^33^.

Interestingly, the evoked potential amplitudes in the auditory cortex during AUD stimulation were reduced compared with those during LCN stimulation. One potential explanation may involve competing interhemispheric effects: direct stimulation of auditory cortex on one side while simultaneously perturbing the contralateral side via cochlear activation. By contrast, during LCN stimulation, cochlear activation of the contralateral Au1 cortex and activation of the DTC pathway represent separate circuits, with limited overlap and reduced interference between them. This distinction may explain why targeting the LCN provides greater leverage than cortical targets in producing pathway-specific responses that can be distinguished from auditory confounds during ultrasound neuromodulation. Importantly, the degree of auditory confounds is likely to vary across tFUS studies depending on the anatomical separation of the stimulation target from cortical networks.

Together, the results support a *convergent model* (Fig. 5C): Cochlear activation by ultrasound-generated sound provides the dominant excitatory drive that reliably activates auditory cortex^34,41^; and tFUS at the target (e.g., LCN) modulates neuronal excitability at subthreshold levels, lowering the activation threshold so that coincident sensory input more effectively recruits downstream circuits. In this framework, cochlear input is necessary to observe robust tFUS-evoked responses in most rodent experiments, but target choice determines pathway bias. The model explains why *Guo* and *Sato et al*^*16,17,32*^ observed little to no effect when cochlear function was eliminated in deaf mouse models -removing the primary trigger^33,34^. While it also aligns with human studies where a somatosensory stimulus paired with tFUS modulated the ERP^11,45^ - the non-auditory trigger replaced the role of sound, and tFUS acted as a gain/threshold modulator for the targeted circuit.

Conceptually, this reframes the “direct vs. indirect” debate: tFUS can do both, but the indirect cochlear drive often dominates the measurable response in rodents, whereas the direct component is best understood as subthreshold excitability modulation that biases specific pathways (here, the DTC) when appropriate input is present. Several biophysical mechanisms have been proposed that are consistent with our convergent model of threshold modulation involving sensory activation. These include brief changes in membrane capacitance (mechanical or flexoelectric)^46,47^, nanoscale cavitation^48^ or modulation of mechanosensitive ion channels (TREK/TRAAK, Piezo, voltage-gated Na^+^/Ca^2+^)^12,46,49,50^, astrocytic TRPA1 signaling^13^, and mild thermal effects^51^. Each of these mechanisms could plausibly modulate neuronal excitability without directly triggering action potentials, providing cellular basis for how tFUS may lower activation thresholds at the target while sensory input—such as cochlear activation—acts as the primary driver of network-level responses.

This places tFUS conceptually alongside other non-invasive brain stimulation techniques, such as tDCS and TMS, where effects primarily arise from shifts in neuronal excitability rather than direct neuronal firing^52,53^. However, unlike these electromagnetic methods, tFUS offers substantially higher spatial precision and depth penetration, enabling selective modulation of deep subcortical targets that remain inaccessible to surface-based stimulation approaches^54^. Practically, this suggests that tFUS could be applied to similar neurological and psychiatric conditions currently treated with tDCS and TMS, such as depression, motor recovery after stroke, or movement disorders^53,55^. Furthermore, rehabilitation of specific circuits may benefit from pairing tFUS with active engagement of the same pathways, either through external sensory inputs (e.g., evoked potentials) or internal voluntary activation (e.g., motor tasks such as hand movement), thereby enhancing synaptic plasticity and promoting targeted functional recovery^56-58^.

One clinical domain where tFUS may demonstrate its greatest potential is in stimulating deep cerebellar nuclei for post-stroke rehabilitation. Modulation of the DTC pathway has already proven effective for promoting motor recovery^21,24,25,59,60^, but conventional approaches rely on invasive deep brain stimulation^22,60,61^. In this context, tFUS offers a distinct advantage— combining non-invasiveness with the ability to reach deep neural structures with millimeter precision. Our finding that LCN-targeted tFUS preferentially modulates contralateral M1 activation suggests a viable strategy for facilitating plasticity within perilesional motor networks. Moreover, if stimulation is combined with task-related paired associative stimuli it could amplify rehabilitation-relevant activity patterns while preserving the spatial selectivity needed to avoid off-target effects.

Our experiments were conducted under anesthesia with modest sample sizes and without direct cochlear blockade; thus, the relative weighting of direct and indirect components cannot be quantified here. Future studies should: (i) manipulate the auditory pathway (acoustic masking, ear occlusion, chemical deafening) and/or use alternative triggers (somatosensory/visual) in deaf models; (ii) map dose–response and parameter regimes that maximize subthreshold modulation while minimizing acoustic artifacts; (iii) record at intermediate nodes (e.g., motor thalamus) to trace signal flow; and (iv) evaluate behavioral outcomes in awake stroke models to establish therapeutic relevance.

Our findings demonstrate that tFUS elicits robust auditory cortex activation consistent with cochlear involvement, alongside a weaker but pathway-specific motor response when deep cerebellar targets are stimulated. These observations support a convergent mechanism in which sensory input provides the principal excitatory drive while tFUS modulates target excitability at subthreshold levels. This framework unifies previously conflicting accounts of tFUS action and offers a principled foundation for harnessing ultrasound neuromodulation in circuit-specific rehabilitation strategies.

## Acknowledgements

This work has been funded by the Science Committee of the Ministry of Science and Higher Education of the Republic of Kazakhstan (Grant No. AP27510389), Nazarbayev University under Faculty-development competitive research grants program for 2025-2027 (Grant No. 040225FD4718) and by the KIST Institutional Program (Project No. 2E33771).

## Author Contributions

A.S. designed and performed the experiments; H.B. and H.K. supported data acquisition and interpretation; A.S. and A.P. analyzed data; D.J. assisted with interpretation of results and figure preparation. A.S. and A.P. wrote the paper with contributions of all coauthors; A.P. supervised research. All authors approved the final version.

