## Supplemental Figure 1 for "Deep cerebellar tFUS engages cortical circuits via convergent local and sensory-driven mechanisms"

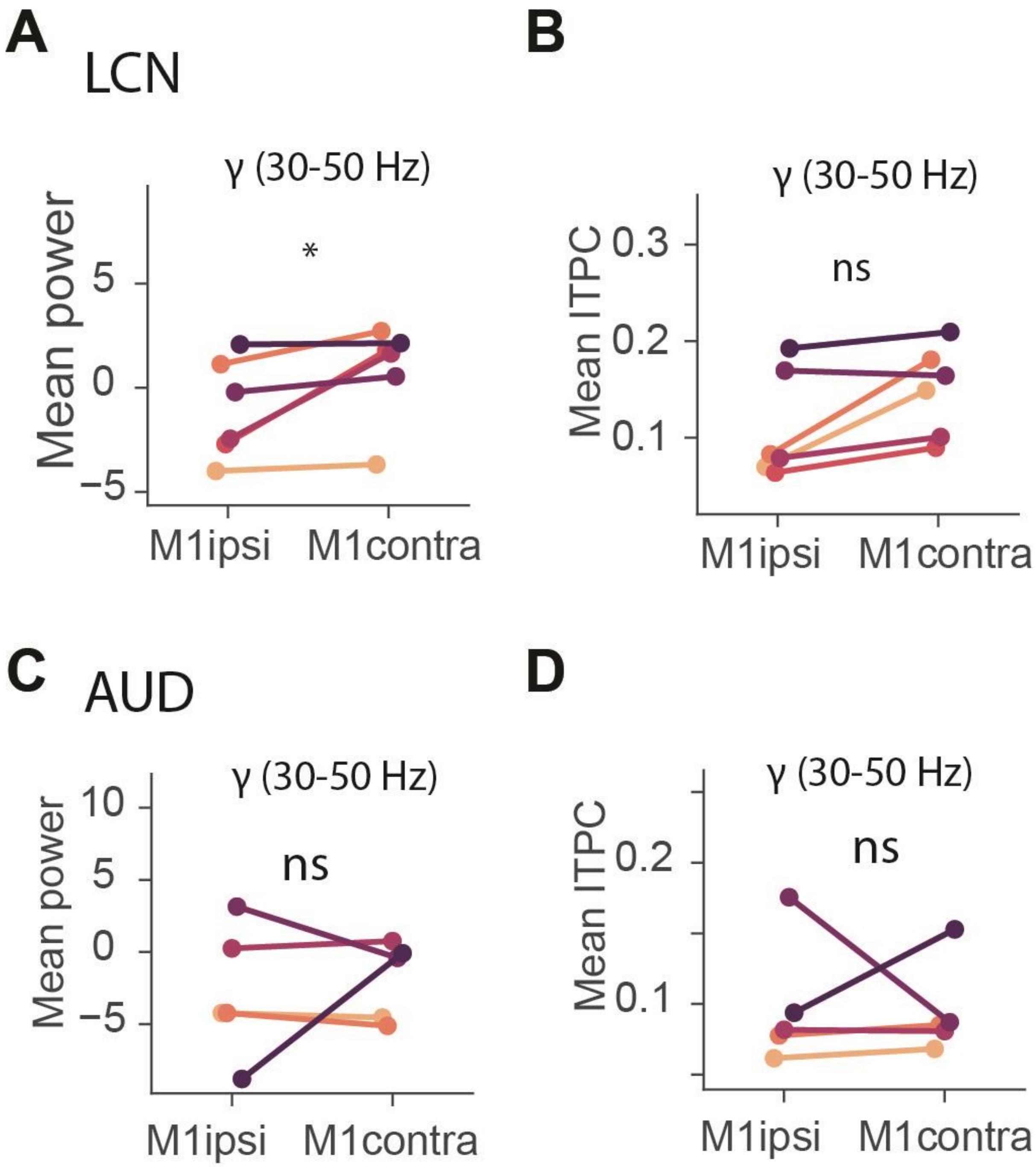


**Figure S1**

A) Quantitative comparison of average spectral power between contralateral and ipsilateral M1 in gamma band (30-50 Hz) within 0-0.2s post-stimulation window for LCN session. Each dot represents one rat (n=6) with connecting lines showing paired measurements (Wilcoxon signed-rank test. *p<0.05).

B) Same as panel B but for ITPC values, quantifying phase consistency across trials within gamma band within 0-0.2s post-stimulation window for LCN session (Wilcoxon signed-rank test. p=0.06).

C) Quantitative comparison of average spectral power between contralateral and ipsilateral M1 in gamma band (30-50 Hz) within 0-0.2s post-stimulation window for AUD session. Each dot represents one rat (n=5) with connecting lines showing paired measurements (Wilcoxon signed-rank test, no significant).

D) Same as panel B but for ITPC values, quantifying phase consistency across trials within gamma band within 0-0.2s post-stimulation window for AUD session. (Wilcoxon signed-rank test, no significant).
